# A Whole Blood Thrombus Mimic: Constitutive Behavior Under Simple Shear

**DOI:** 10.1101/2020.07.19.210732

**Authors:** Gabriella P. Sugerman, Sotirios Kakaletsis, Parin Thakkar, Armaan Chokshi, Sapun H. Parekh, Manuel K. Rausch

## Abstract

Deep vein thrombosis and pulmonary embolism affect 300,000-600,000 patients each year in the US. The progression from deep vein thrombus to pulmonary embolism occurs when blood clots, as a whole or partially, break off from the deep veins and eventually occlude the pulmonary arteries. Venous thromboembolism is the cause of up to 100,000 deaths per year in the US alone. To date, we don’t fully understand this mechanical process among other reasons because in-vivo samples are difficult to obtain, highly heterogeneous, and their shapes are inappropriate for most mechanical tests. Toward overcoming these obstacles, we have set out to develop an in-vitro thrombus mimic and to test this mimic under large deformation simple shear. In addition to reporting on the mechanics of our mimics under simple shear, we explore the sensitivity of their mechanics to coagulation conditions and blood storage time, and compare three hyperelastic material models for their ability to fit our data. We found that thrombus mimics made from whole blood demonstrate strain-stiffening, a negative Poynting effect, and hysteresis when tested quasi-statically to 50% strain under simple shear. Additionally, we found that the stiffness of these mimics does not significantly vary with coagulation conditions or blood storage times. Of the three hyperelastic constitutive models that we tested, the Ogden model provided the best fits to both shear stress and normal stress. In conclusion, we developed a robust protocol to generate regularly-shaped, homogeneous thrombus mimics that lend themselves to simple shear testing under large deformation. Future studies will extend our model to include the effect of maturation and explore its fracture properties toward a better understanding of embolization.

## 1. Introduction

Deep vein thrombosis and pulmonary embolism affect 300,000-600,000 patients each year in the US alone (Cushman, 2007). Collectively known as venous thromboembolism, these disorders are characterized by pathologic blood coagulation restricting or eliminating blood flow through the veins (Kearon, 2003). The initial thrombus is subject to a variety of possible outcomes: it may resolve with relatively few symptoms (Dewyer et al., 2007; Nguyen et al., 2015), persist and reduce blood flow causing post-thrombotic syndrome (Ashrani and Heit, 2009), or break apart and shed emboli into the bloodstream, which in turn may result in obstruction of flow to critical organs, such as the lung (van Langevelde et al., 2013). The latter outcome is the most catastrophic leading to approximately 100,000 deaths every year in the US alone according to the CDC (Beckman et al., 2010). The deformation, fracture, and eventual embolization of thrombus are inherently mechanical processes (Basmadjian, 1989; Johnson et al., 2017). Thus, the mechanics of thrombus, and venous thrombus in particular, has been of immense interest.

Investigations of thrombus mechanics have taken many forms. Because fibrin is the primary structural constituent of early thrombi, there has been intense effort invested in studying and characterizing the mechanics of fibrin networks (Piechocka et al., 2010; Brown et al., 2009; Liu et al., 2006; Fleissner et al., 2016). However, in reality, thrombi are composed of not only fibrin, but also so called formed elements, platelets and red blood cells. Thus, other studies have also considered the role of cellularity in thrombus mechanics (Gersh et al., 2009; Kim et al., 2017). These studies have found that closely-packed elements significantly alter the mechanical behavior of thrombus (Tutwiler et al., 2018; van Oosten et al., 2019) requiring a holistic approach toward studying thrombus. Additionally, the constitution and thus mechanics of in-vivo thrombi are not time-invariant (Rausch and Humphrey, 2017). For example, we have previously probed the uniaxial tensile behavior of venous thrombi developed in mice at two time points: after two weeks and after four weeks of formation (Lee et al., 2015). Additionally, we characterize the thrombi’s constitution via histology. We found that thrombus “maturation” (i.e., resident time in-vivo) profoundly influenced the compositional and mechanical properties of the explanted thrombi. Hence, to develop a fundamental understanding for venous thrombus mechanics we must consider thrombi in their full complexity as well as across time scales. While animal models for deep vein thrombus can be useful tools, as shown in our previous study, they provide significant challenges in addition to costing animal lives. For example, thrombus maturation is not a homogeneous process. We found that thrombi are invaded with immune and synthetic cells that begin converting the initial fibrin-based thrombus into a collageneous pseudo-tissue from the radial boundary inward. The arising spatial heterogeneity of thrombus renders these tissue complex model systems. Additionally, in-vivo thrombus are not regularly-shaped and make testing via standard methods difficult.

To study the mechanics of venous thrombus while overcoming the shortcomings of in-vivo studies, we aim to develop an in-vitro model of evolving venous thrombus. In our current work, we established the methodology for generating venous thrombus mimics based on fresh coagulated blood and investigated their baseline mechanical behavior. To this end, we developed a simple shear testing protocol, applied our test protocol to thrombus mimics made from bovine blood, and reported the material parameters for three hyperelastic material models. Additionally, we tested the sensitivity of these mimics’ mechanics to coagulation conditions. Specifically, we varied both the amount of calcium chloride used to reverse our anticoagulant and the coagulation time on the shear deformation behavior, and report on the effect of blood storage time.

## 2. Methods

### 2.1. Thrombus mimic generation

We generated thombi from bovine blood (Lampire Biological Laboratories, PA, USA) that was collected with CPDA-1 anticoagulant at 14% volume:volume and stored at 4 °C for 12-72 hours prior to experimentation. We initiated coagulation by adding calcium chloride to a final concentration of 10, 20, or 40 mM (Roessler et al., 2014; Kim et al., 2017). During coagulation, we gently mixed blood with a wide-mouth pipette, and then injected it into 12×10×10 mm cube molds lined on two opposing sides with hook-and-loop fabric such that the cube volume was 1000 mm^3^, see Figure 1a. Samples coagulated for 60 min, 90min, or 120 min at 37 °C before we removed them from the mold, see Figure 1b. We defined the start of coagulation time when calcium chloride was added, while we defined the end of coagulation time when shear testing experiments commenced.

**Figure 1:**
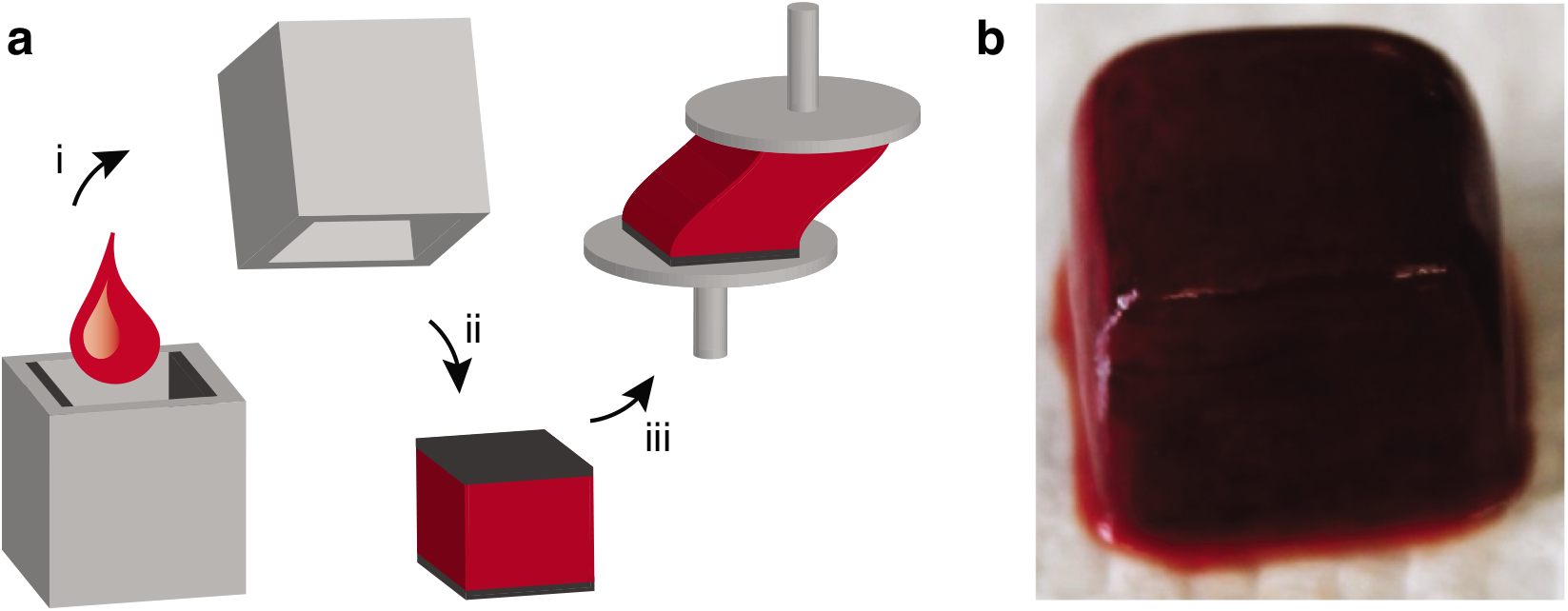
Generation of thrombus mimics. (a) (i) We mixed anticoagulated whole bovine blood with calcium chloride and coagulated both in cube-shaped molds lined on two opposing sides with hook-and-loop fabric. (ii) After coagulation, we removed samples with integrated hook-and-loop fabric from molds. (iii) We secured the mimics to pin stubs with a drop of super glue on the back of the hook-and-loop fabric. (b) Exemplary thrombus mimic (shown without hook-and-loop fabric to illustrate sample quality.)

### 2.2. Shear testing

Following the specified coagulation time, we removed samples from their molds and secured the hook-and-loop fabric-covered ends to pin stubs via super glue. Note, the super glue bonded the smooth surfaces of the hook-and-loop fabric to the pin stub. In other words, the super glue did not contact the thrombus material at any point. We let the glue dry for one minute before we submerged the samples in phosphate-buffered saline (PBS) at 37 °C ±2. Throughout testing, we maintained sample temperature by circulating attemperature PBS through our bath.

We conducted simple shear experiments by symmetrically displacing the top plate of our custom simple shear testing device by ±5 mm in each direction at a rate of 0.1 mm/s for ten consecutive cycles. Simultaneously, we measured force in three directions using a 2N capacity triaxial load cell. With a custom LabView program, we collected time, displacement, and force in shear and normal directions during testing.

### 2.3. Inverse Analysis

Simple shear is not so simple (Destrade et al., 2012). It is well-known that non-zero tractions on the incline surface may be required to maintain a homogeneous deformation field during simple shear experiments (Horgan and Murphy, 2011). Given that simple shear testing apparati maintain traction free conditions on the incline surface, homogeneous solutions to simple shear are likely inaccurate. Schmid et al. have previously demonstrated the need for inverse finite element analysis to extract material parameters for soft tissue from simple shear experiments (Schmid et al., 2008). Thus, we too developed an (iterative) inverse finite element analysis framework using Matlab R2019a and FEBio 2.9.1 to identify the material parameters for our thrombus mimics (Maas et al., 2012).

Specifically, for the forward simulations, we modeled each sample as a cube with 10 mm edge length discretized by an equal number of linear hexahedral mixed elements along each edge. We defined the boundary conditions by fixing displacements of the sample’s bottom surface, while attaching the top surface nodes to a rigid body. In turn, we constrained the rigid body to displace in only the shear direction. During the simulation, we applied the experimental pin stub displacement profile to the rigid body, while measuring its reaction forces to the deforming sample. See Figure 2 for a sample simulation which depicts the highly heterogeneous deformation field under simple shear.

**Figure 2:**
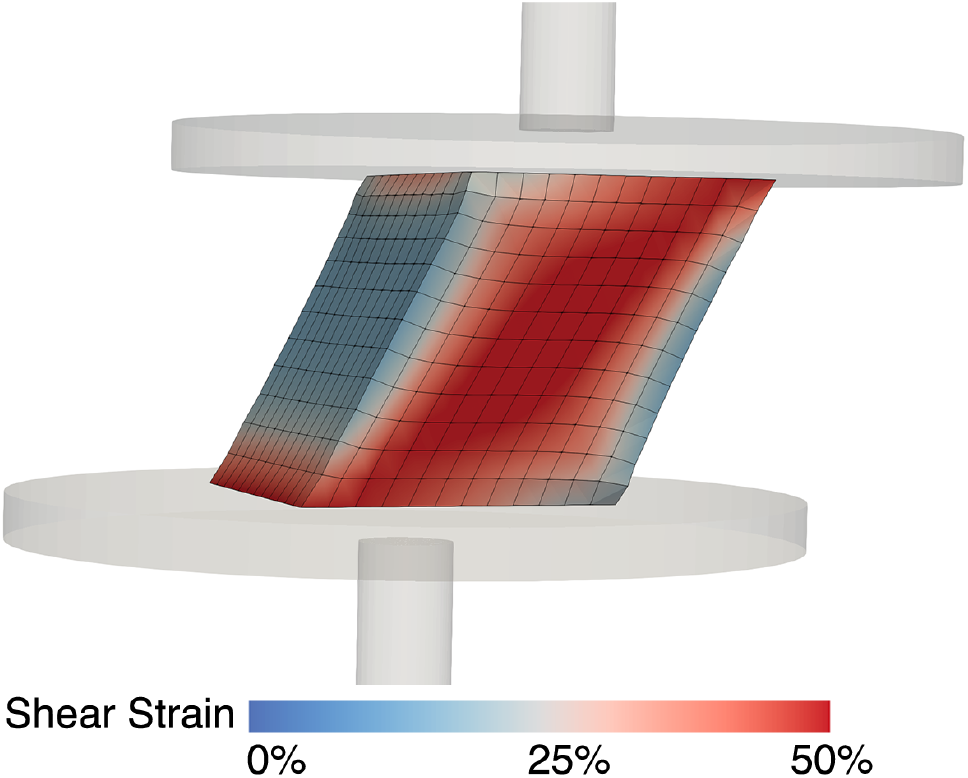
Simple shear is not so simple. Locally resolved Green-Lagrange shear strain *γ* = 2*E_xz_* derived from our inverse finite element simulation, where *E_xz_* is the off-diagonal shear element of the Green-Lagrange strain tensor.

For the constitutive fitting, we tested three hyperelastic material models, assumed to be nearly incompressible. To this end, we made use of the decoupled material formulation in FEBio that decomposes the deformation gradient **F** with the invariant *J* = *det*(**F**) into an isochoric and volumetric part, i.e., 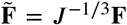 and *J*^1/3^**I**, respectively. Accordingly, the deviatoric right Cauchy-Green tensor reads 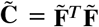, while the deviatoric principal stretches read 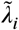 and the first deviatoric invariant becomes 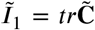 (Holzapfel, 2000). Furthermore, *U*(*J*) is the volumetric component of the strain energy, for which FEBio assumes the form 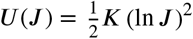 with bulk modulus *K*. Specifically, we identified the parameters to the following models:

1. Ogden model (Ogden, 1973)

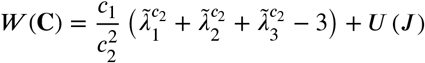
2. Isotropic Fung-type (Fung, 1967; Demiray, 1972; Holzapfel et al., 2000)

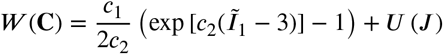
3. Yeoh model (Yeoh, 1993)

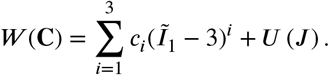

Following a convergence analysis for all three models on the peak tangent and normal force values, we performed all simulations using a 12^3^-element cube. For all tested samples, we estimated the values of the parameter vector **c** using non-linear least squares regression in Matlab. To this end, we defined the objective function *Z*(**c**) to be minimized as follows

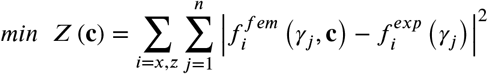

where *x* and *z* imply the shear and normal force, respectively, over *n* =100 shear strain points *γ_j_*. Thus, we fitted both the shear force and the normal force simultaneously. Specifically, we used Matlab to call FEBio with an initial guess for the material model parameters, performed a forward simulation in FEBio, read the simulation-based output forces in Matlab, computed the error metric via above objective function, and repeated this process iteratively until reaching our error minimum. Finally, for the optimal set of parameters **c**\*, we evaluated goodness of fit across samples by comparing the normalized mean-square error (NMSE)

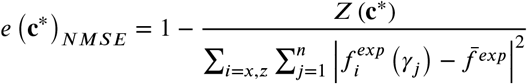

where 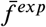 is the mean recorded force over all data points for both normal and shear forces.

### 2.4. Histology

We generated thrombus mimics specifically for histology using 20 mM calcium chloride and a coagulation time of 60 minutes. We then fixed those samples in 10% neutral buffered formalin for 48 hours. Subsequently, we dehydrated the samples in ethanol, embedded them in paraffin, and sectioned them at 5 μm thickness. Following standard protocols, we stained all slides with hematoxylin and eosin (H&E) to show red and white blood cells. We acquired histological images at 4, 10, 20 and 40× magnification with a BX53 (Olympus, Tokyo, Japan) microscope and an DP80 camera (Olympus, Tokyo, Japan) using bright-field imaging.

### 2.5. Cell Counting

We monitored viability of red blood cells and platelets by isolating and counting cells from blood stored for 1,2, and 3 days. To this end, we centrifuged blood for 30 minutes at 200×g to separate platelet-rich plasma (PRP) from red blood cells. We mixed PRP with an equal volume of HEPES buffer (140 mM NaCl, 2.7 mM KCl, 3.8 mM HEPES, 5 mM EGTA, in ultra-pure water) with prostaglandin E1 (1 μM final concentration) then centrifuged the solution for 15 minutes at 100×g to pellet contaminating cells and transferring PRP to a new tube. Finally, we centrifuged the PRP once more at 800×g to pellet platelets which we subsequently washed in EDTA buffer (10 mM sodium citrate, 150 mM NaCl, 1 mM EDTA, 1% (w/v) dextrose) and resuspended in modified Tyrode’s buffer (Tyrode’s (HIMEDIA), 5 mM dextrose, 3 mg/mL bovine serum albumin, 1 μM prostaglandin E1). Separately, we resuspended the red blood cell layer win 0.9% NaCl (aqueous) and centrifuged it for 5 minutes at 500×g. We performed this step two additional times to wash and pack the red blood cells. Finally, we suspended the red blood cells in a storage buffer (150 mM NaCl, 5mM dextrose, 3 mg/mL bovine serum albumin). Before counting red blood cells and platelets in a hemocytometer, we combined each solution of isolated cells with Trypan Blue (0.4% aqueous) in a 1:4 ratio.

### 2.6. Statistical analysis

We examined differences in toe-region stiffness and calf-region stiffness as functions of calcium chloride concentration and coagulation time using a Friedman test. Importantly, given our replicate number of three per group, our calf-stiffness correlation and variation, expectation of a large effect (Cohen’s for-many-means-effect size of 0.4), and correction for the use of the Friedman test via its asymptotic relative efficiency, we reach a power 0.84. We performed the actual power analysis in G*Power (Version 3.1). Furthermore, we tested the time dependence of toe-region and calf-region stiffness as well as red blood cell and platelet count via least squares regression and Spearman correlation. We performed our statistical analyses in MATLAB R2019a. We considered results as significantly different for p-values smaller than 0.05. Throughout this text, we present data as mean ±1 standard deviation.

## 3. Results

Coagulation in molds yielded cubes of thrombus mimics which were ideal for histological analyses and simple shear experiments.

First, to illustrate the mimics’ microstructure and composition, we stained samples via H&E. Figure 3 shows H&E stains of a typical mimic at four magnification, from 4× to 40×. H&E identifies white blood cell nuclei in dark purple and red blood cells in pink. Thus, it demonstrates the relative abundance of red blood cells compared to white blood cells. Under lower-magnification (i.e., 4× and 10×) the sample appears mostly homogeneous with some areas of higher density, while higher-magnification images show cell-level inhomogeneity (i.e., 20× and 40×). We interpret white space as areas of low cell density which are likely composed of fibrin network, as fibrin is not stained by H&E. Note, samples did not demonstrate gradients in red blood cell density, which would indicate settling of cells during coagulation, as reported by others (Chernysh et al., 2020). Overall, our thrombus mimics appeared homogeneous without structural elements that would render it anisotropic.

**Figure 3:**
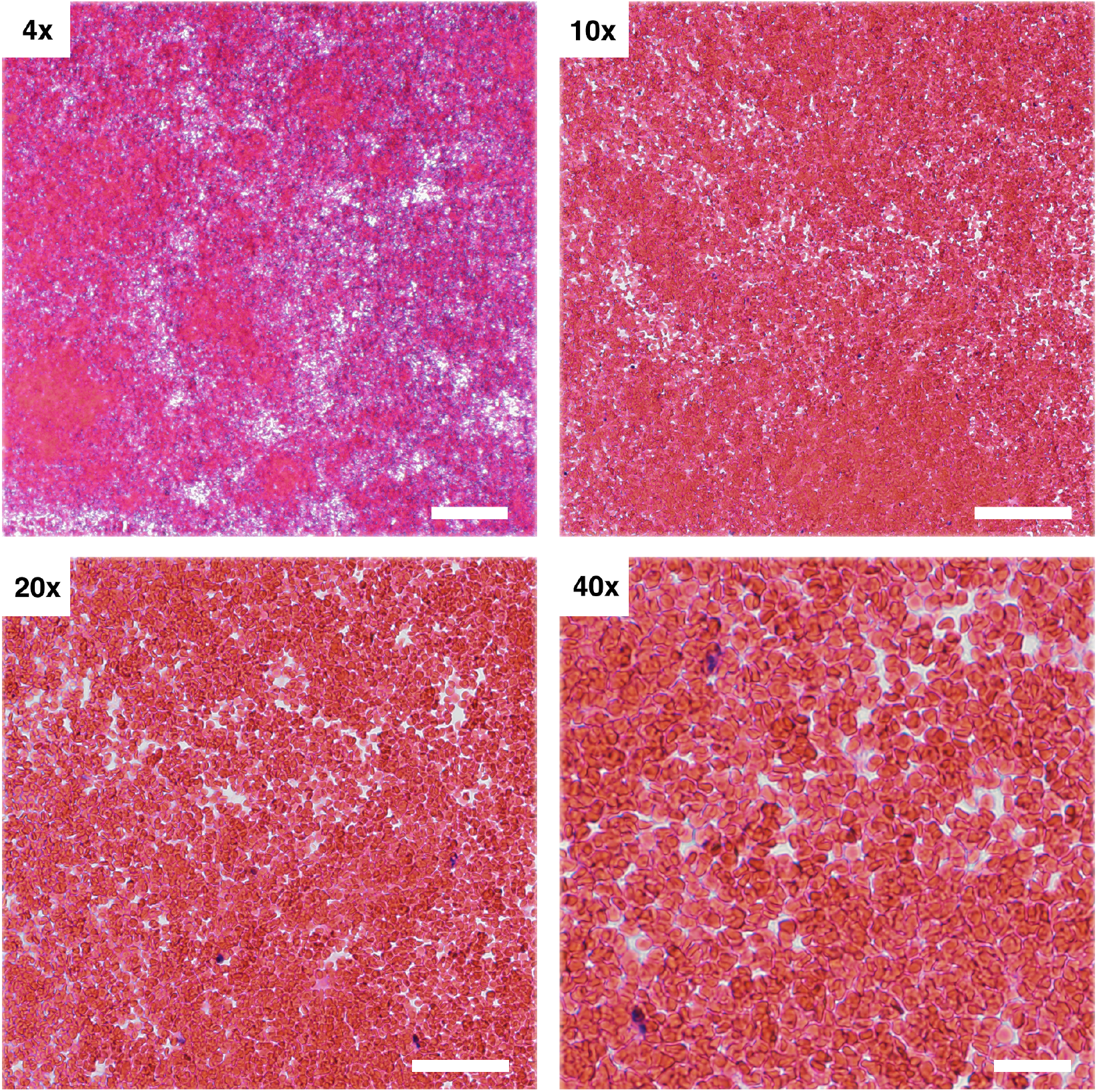
Thrombus mimics were predominantly homogeneous without a discernible density gradient. We took images of H&E-stained thrombus mimics at 4X, 10X, 20X, and 40X magnification. Our scale bars are 200, 100, 50, and 20 μm, respectively. Lower-magnification images (i.e., 4X and 10X) appear mostly homogeneous with some areas of higher density, while higher-magnification images show cell-level inhomogeneity (i.e., 20X and 40X). The shown thrombus mimic was coagulated with 20 mM calcium chloride for 60 min.

Next, our primary goal was to investigate the constitutive behavior of our thrombus mimic via simple shear experiments. To this end, Figure 4a depicts the shear stress of a representative mimic under simple shear. In complement, Figure 4b shows the normal stress of the same sample. We found that samples under large deformation showed the classic “J-shaped” strain-stiffening behavior known from many other fibrous soft tissues (Rausch and Humphrey, 2016; Meador et al., 2020b,a). Samples also demonstrated some hysteresis, i.e., difference between loading and unloading. Interestingly, we found that our samples exhibited negative normal stress. In other words, when sheared our samples contracted rather than expanded, i.e., exhibited a negative Poynting effect (Mihai and Goriely, 2011; Janmey et al., 2007). In the subsequent text, when reporting on both shear stress and normal stress, we refer to the average (or midline) of the upstroke and downstroke, ignoring the material’s viscoelasticity in a first approximation. Furthermore, to compare the mechanical behavior of thrombus mimics across various coagulation conditions, we extracted scalar metrics from these nonlinear stress-strain curves: first, we computed the tangent modulus at low strains (i.e., toe-stiffness). Second, we computed the tangent modulus at high strains (i.e., calf-stiffness), see Figure 4c. In summary, under simple shear our mimics exhibited a non-linear strain-stiffening response to shear while demonstrating a negative Poynting effect and hysteresis.

**Figure 4:**
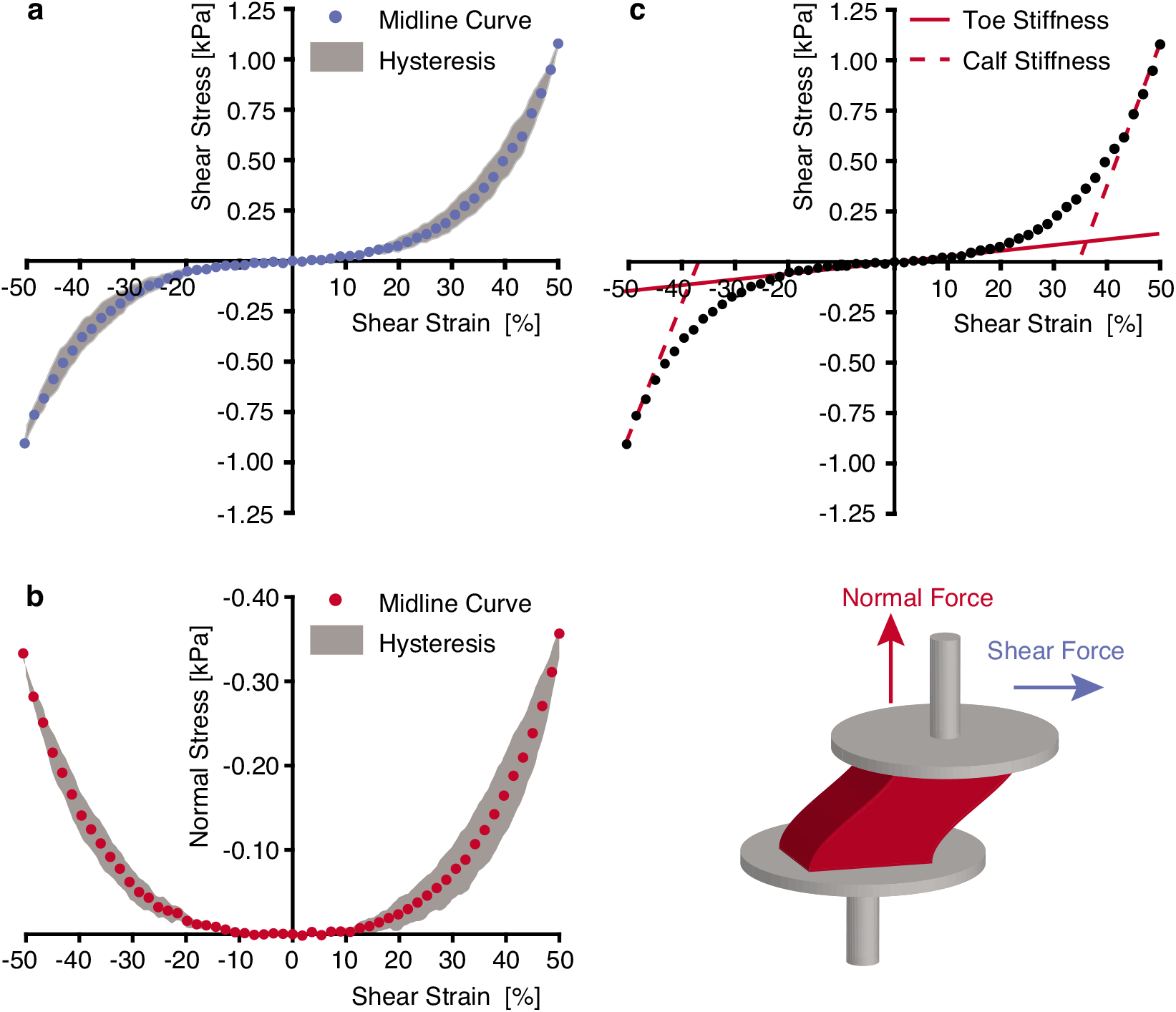
Thrombus mimics demonstrated shear-stiffening, negative normal stress, and hysteresis. Stress-strain behavior of a representative sample under simple shear: (a) Shear stress and (b) normal stress after nine preconditioning cycles. As a first approximation, we subsequently represent the mechanical behavior of our thrombus mimics by the average midline of the upstroke and downstroke (circles). (c) Depiction of tangent modulus at low strain (toe-stiffness) and at high strain (calf-stiffness). This representative thrombus mimic was coagulated with 20 mM calcium chloride for 60 min.

To explore the sensitivity of our thrombus mimics to coagulation conditions, we performed simple shear experiments on samples coagulated under various conditions. Figure 5 shows the average shear stress (± 1 standard deviation) across three samples for each coagulation time and calcium concentration. Qualitatively all curves look consistent across changes in calcium chloride concentration from 10 to 40 mM and coagulation times from 60 to 120 minutes. We also quantitatively explored the sensitivity of our thrombus mimics to coagulation times and calcium chloride concentration by comparing toe- and calf-stiffness between samples. Figure 6a illustrates the insensitivity of toe-stiffness to changes of coagulation time or calcium chloride concentration (time: p = 0.6605, concentration: p = 0.3292). Similarly, Figure 6b shows that clots with varying coagulation characteristics maintained similar stiffness in the calf region (time: p = 0.2174, concentration: p = 0.3244). Through qualitative and quantitative comparisons, we didn’t find a significant effect of coagulation conditions.

**Figure 5:**
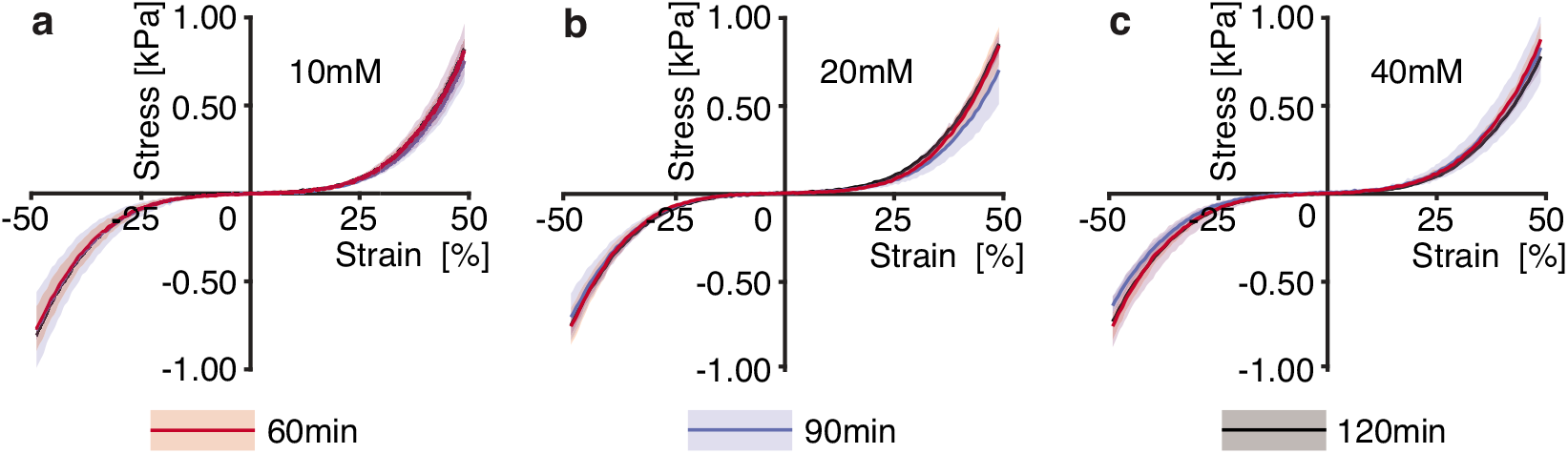
Shear stress appeared insensitive to changes in coagulation time or calcium chloride concentration. Mean and standard deviation of midline shear stress for samples coagulated for 60, 90, and 120 minutes with (a) 10 mM, (b) 20 mM, (c) 40 mM calcium chloride. Each line represents an average across three samples.

**Figure 6:**
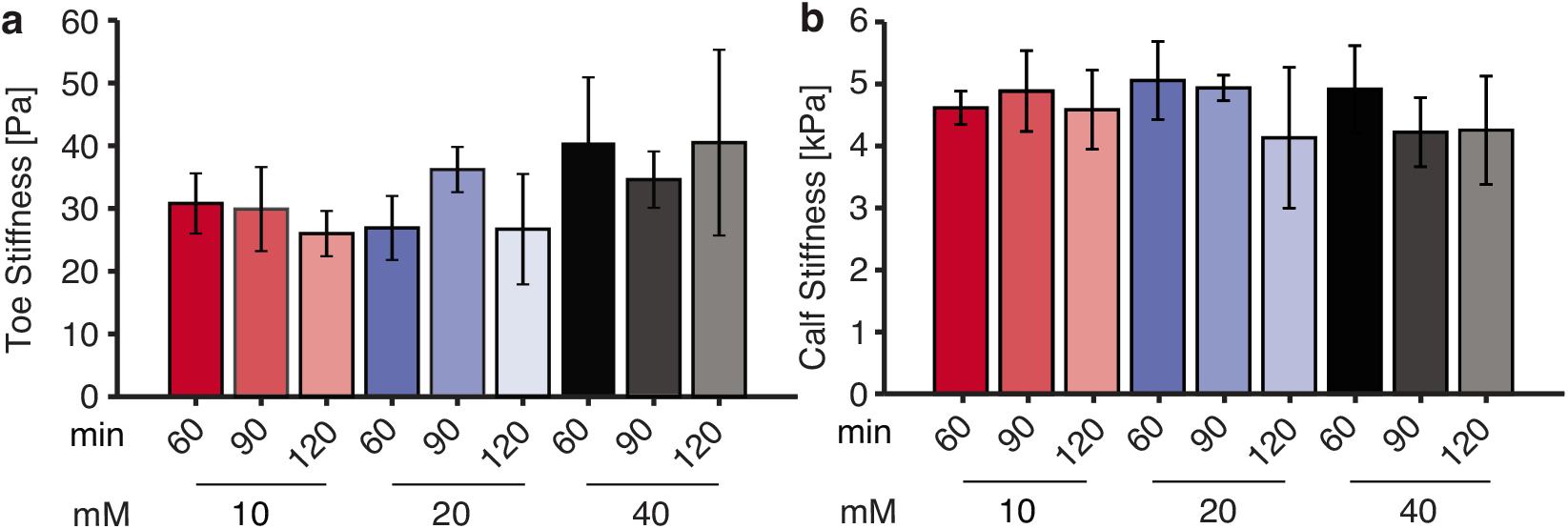
Thrombus mimic stiffness didn’t appear to depend on calcium chloride concentration or coagulation time. Tangent stiffness computed at (a) low strain (toe-stiffness, n=3 per group), (b) high strain (calf-stiffness, n=3 per group).

Because we tested samples after 24 to 72 hours of storage (1-3 days), we were concerned that cell viability may affect our results. To this end, we report red blood cell and platelet count over time, see Figure 7a-b. Additionally, we compared shear stiffness metrics as a function of storage time, see Figure 7c-d. Least squares fits to all data declined marginally over three days, indicated by the negative R-values. However, none of the Spearman correlations were significant (p>0.05). Thus, it appears that storage times between 24 and 72 hours did not affect mechanical properties of our thrombus mimics.

Lastly, to identify the material parameters to our thrombus mimics, we fit our midline shear and normal force data using the Ogden, Yeoh, and Fung-type constitutive models.

**Figure 7:**
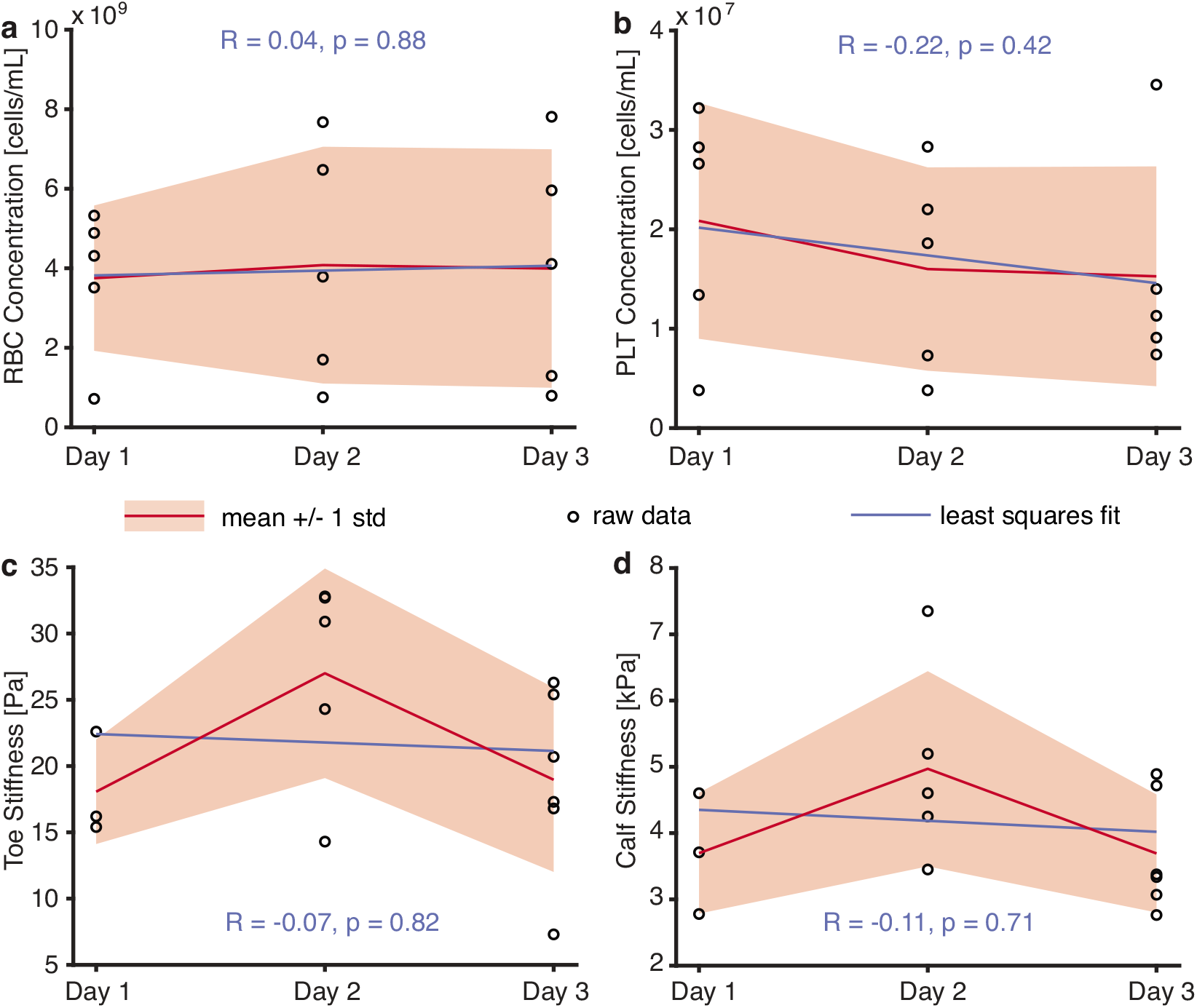
Cell counts and stiffness of thrombus mimics didn’t appear to depend on storage time between 24-72 hours. Manual counts of (a) red blood cells, (b) platelets (n=5 counts per day). (c) Toe-stiffness and (d) calf-stiffness of thrombus mimics coagulated with 20mM calcium chloride for 60min (n=3,4,5 per group, by day, respectively).

Figures 8a-c depict fits of all models to shear and normal stress for the same sample data set that was also depicted in Figure 4. The goodness-of-fit metric normalized mean-squared error (NMSE) is inset in each subfigure. Note, a perfect fits yields a NMSE of 1. Thus, the higher the NMSE the better the fit. See Table 1 for details on all tested samples, parameters, and goodness-of-fit data. Overall, of the three models the Ogden model fit both the shear data as well as the normal data qualitatively and quantitatively better than the other two models.

**Figure 8:**
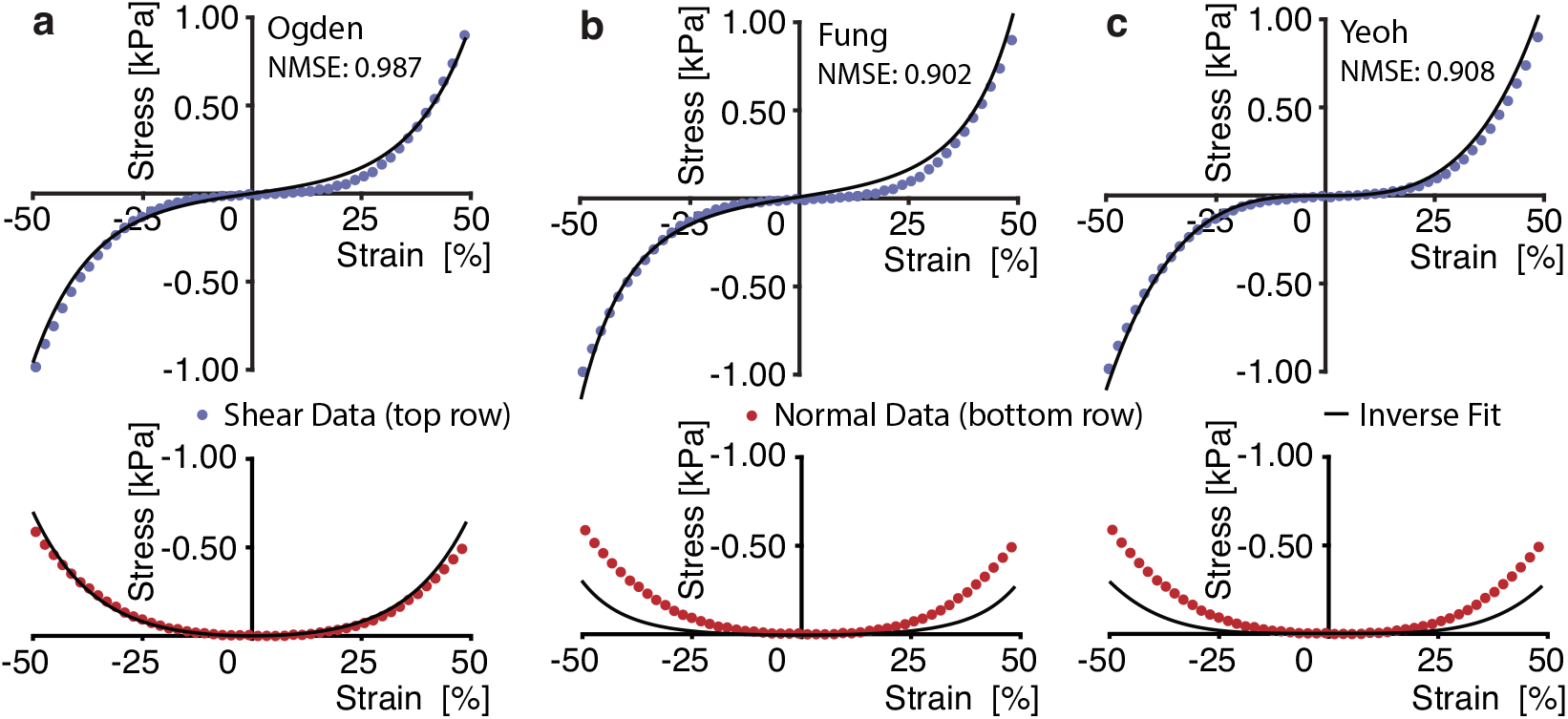
The Ogden model fit the mechanical behavior of our thrombus mimics best. Representative inverse model fits of shear (blue, top) and normal (red, bottom) stress for (a) Ogden, (b) Fung, (c) Yeoh models. Also shown are normalized mean-square error (NMSE) for each fit. This representative thrombus mimic was coagulated with 20mM calcium chloride for 60min. Note, a perfect fits yields a NMSE of 1. Thus, the higher the NMSE the better the fit.

**Table 1.** Estimated material parameters of all 3 constitutive models, for every sample tested, with corresponding normalized mean squared errors (NMSE) where A: Concentration of calcium chloride [mM], B: Coagulation time [min], and C: Replicate Number. Note, a perfect fits yields a NMSE of 1. Thus, the higher the NMSE the better the fit.

| A | B | C | Ogden Parameters |  | Fung Parameters |  | Yeoh Parameters |  |  | NMSE |  |  |
| --- | --- | --- | --- | --- | --- | --- | --- | --- | --- | --- | --- | --- |
| | | | $c_1$ (Pa) | $c_2$ (-) | $c_1$ (Pa) | $c_2$ (-) | $c_1$ (Pa) | $c_2$ (Pa) | $c_3$ (Pa) | Ogden | Fung | Yeoh |
| 10 | 60 | 1 | 584.28 | 16.35 | 329.21 | 8.54 | 0.25 | 1655.26 | 2296.56 | 0.985 | 0.893 | 0.899 |
|  |  | 2 | 851.56 | 15.29 | 453.32 | 8.00 | 12.79 | 2324.01 | 1751.37 | 0.985 | 0.910 | 0.915 |
|  |  | 3 | 768.91 | 15.65 | 412.01 | 8.29 | 1.60 | 2126.79 | 2189.50 | 0.979 | 0.919 | 0.924 |
|  | 90 | 1 | 622.05 | 16.22 | 355.91 | 8.35 | 18.84 | 1613.72 | 2642.16 | 0.984 | 0.909 | 0.914 |
|  |  | 2 | 772.41 | 15.91 | 419.93 | 8.41 | 0.01 | 2189.79 | 2388.77 | 0.983 | 0.911 | 0.917 |
|  |  | 3 | 819.98 | 15.55 | 434.81 | 8.27 | 21.72 | 2113.08 | 2493.28 | 0.984 | 0.918 | 0.923 |
|  | 120 | 1 | 530.39 | 16.32 | 300.75 | 8.47 | 21.69 | 1381.59 | 2209.34 | 0.989 | 0.890 | 0.895 |
|  |  | 2 | 611.11 | 15.99 | 305.47 | 9.14 | 8.97 | 1409.62 | 3390.48 | 0.956 | 0.955 | 0.960 |
|  |  | 3 | 961.40 | 15.35 | 526.70 | 7.83 | 7.06 | 2741.11 | 1672.24 | 0.987 | 0.902 | 0.908 |
| 20 | 60 | 1 | 560.82 | 16.38 | 298.93 | 9.01 | 1.54 | 1456.56 | 2922.19 | 0.972 | 0.931 | 0.937 |
|  |  | 2 | 837.06 | 15.64 | 431.86 | 8.57 | 0.08 | 2204.81 | 2935.81 | 0.973 | 0.933 | 0.939 |
|  |  | 3 | 650.69 | 16.26 | 351.59 | 8.81 | 0.13 | 1773.09 | 2873.17 | 0.976 | 0.925 | 0.932 |
|  | 90 | 1 | 777.55 | 15.64 | 412.33 | 8.34 | 0.08 | 2163.39 | 2157.83 | 0.981 | 0.916 | 0.922 |
|  |  | 2 | 717.81 | 15.46 | 384.11 | 8.10 | 4.12 | 2010.73 | 1579.70 | 0.983 | 0.906 | 0.912 |
|  |  | 3 | 899.49 | 15.14 | 490.33 | 7.66 | 15.36 | 2536.95 | 1132.92 | 0.990 | 0.889 | 0.895 |
|  | 120 | 1 | 515.83 | 15.15 | 266.30 | 8.13 | 20.92 | 1199.10 | 1655.79 | 0.971 | 0.935 | 0.940 |
|  |  | 2 | 621.25 | 16.09 | 316.57 | 9.06 | 0.16 | 1555.22 | 3140.16 | 0.965 | 0.944 | 0.950 |
|  |  | 3 | 847.24 | 15.38 | 458.64 | 7.94 | 11.78 | 2378.30 | 1564.58 | 0.988 | 0.898 | 0.904 |
| 40 | 60 | 1 | 747.15 | 15.07 | 381.16 | 8.09 | 2.75 | 1961.91 | 1718.34 | 0.975 | 0.923 | 0.930 |
|  |  | 2 | 1401.10 | 11.23 | 487.07 | 7.54 | 100.80 | 1775.62 | 2658.05 | 0.927 | 0.988 | 0.991 |
|  |  | 3 | 890.52 | 14.68 | 446.89 | 7.90 | 2.98 | 2404.48 | 1241.52 | 0.977 | 0.932 | 0.938 |
|  | 90 | 1 | 657.78 | 16.17 | 361.92 | 8.58 | 0.03 | 1837.67 | 2541.12 | 0.981 | 0.914 | 0.921 |
|  |  | 2 | 799.69 | 13.87 | 374.07 | 7.67 | 30.06 | 1764.43 | 1296.11 | 0.970 | 0.946 | 0.951 |
|  |  | 3 | 1054.73 | 14.18 | 580.77 | 6.75 | 55.58 | 2742.89 | 0.04 | 0.990 | 0.878 | 0.884 |
|  | 120 | 1 | 593.99 | 15.37 | 319.31 | 8.01 | 8.07 | 1651.90 | 1191.73 | 0.982 | 0.907 | 0.912 |
|  |  | 2 | 898.52 | 14.22 | 457.24 | 7.38 | 0.10 | 2333.46 | 988.79 | 0.971 | 0.932 | 0.940 |
|  |  | 3 | 1090.10 | 13.39 | 536.76 | 6.90 | 26.29 | 2695.45 | 0.05 | 0.979 | 0.925 | 0.932 |

## 4. Discussion

### 4.1. Summary

We have successfully developed an in-vitro approach that yields regularly-shaped thrombus mimics that lend themselves well to simple shear testing under large deformation. This methodology is a first step on our path toward a robust in-vitro model for venous thrombus mechanics. In our thrombus mimics, we measured shear and normal stress and compared metrics of shear stiffness as a function of various coagulation conditions including calcium chloride concentration, coagulation times, and blood storage time.

### 4.2. Why simple shear?

We chose simple shear as the mode of deformation for two reasons: First, shear is likely thrombus’ predominant in-vivo mode of deformation. Therefore, our testing modality reflects the natural mode of deformation most accurately. Second, simple shear is approximately volume preserving (Gardiner and Weiss, 2001). We believe that this minimizes the effects of intra-sample pressure gradients that lead to porous fluid flow and thus energy-dissipating momentum exchange between the fluid phase and the solid phase of the clearly poroelastic material (Suh and DiSilvestro, 1999). In turn, such theoretical decoupling reduces the overall viscoelastic phenomena and improves our approximation of mimic material as hyperelastic.

### 4.3. Mechanics of our thrombus mimic

Qualitatively, our mechanical testing via simple shear revealed a strain-stiffening material with negative Poynting effect and hysteresis. As to the strain-stiffening of thrombus, past reports have been mixed. While some studies showed that thrombus is strain-stiffening (Lee et al., 2015; Teng et al., 2015) others showed that it behaves linearly (Di Martino et al., 1998; Gasser et al., 2008; Geest et al., 2006). Disagreement between data likely stems from the drastic variation between samples due to differences in coagulation conditions (under flow versus stasis, on the arterial side versus venous side, pre-mortem versus post-mortem, in-vitro versus in-vivo, etc.). Additionally, thrombus testing methods have varied, employing uniaxial tensile and compressive testing, planar biaxial testing, nanoindendation, small and large angle shear rheometry, and now simple shear (this publication). For a comprehensive review of the thrombus mechanics literature please see Johnson et al. and the works cited therein (Johnson et al., 2017). Interestingly, van Oosten et al. have argued that it is the densely-packed cellular elements in thrombus that may render it non-shear stiffening. Disagreement between their argument and a number of publication demonstrating otherwise have so far not been discussed (to the best of our knowledge). As to our finding that thrombus exhibits a negative Poynting effect: This means that our thrombus mimics pulled on our pin stubs while they were sheared rather than trying to expand them. In contrast, other soft tissues such as brain show the opposite effect (Destrade et al., 2012). This effect has recently been attributed to the strain-stiffening response of the underlying fibrin network (Janmey et al., 2007) and has further been discussed in the context of rubber-theory (Mihai and Goriely, 2011). Additionally, it has been proposed that it is the clot porosity that may give rise to the negative Poynting effect seen in our data (de Cagny et al., 2016). Finally, all soft tissues exhibit viscoelastic behavior as indicated by differences between the loading and unloading curve (i.e., hysteresis). As mentioned above, we chose simple shear as our test modality to minimize viscoelastic effects in our data (Gardiner and Weiss, 2001). However, while the majority of viscoelasticity in thrombus likely arises from its poroelasticity, some unarguable also arises from the solid-phase viscoelasticity of fibrin (Rausch et al., 2009). Future studies may explore the relative contribution between these two sources in a dedicated study.

Quantitatively, we have few studies to compare our data to as there have been no large deformation simple shear experiments on in-vitro thrombus mimics before. However, we can compare our findings to the findings of others as we reduced the constitutive response of our mimics to two scalar metrics, i.e., toe-stiffness and calf-stiffness, which represent the full range of our mimics’ stiffness. Specifically, we found toe-stiffness of ≈ 30Pa and the calf-stiffness of≈ 5kPa. These values indicate that thrombus is “ultra-soft” and compare well with findings of others (Riha et al., 1999).

### 4.4. Constitutive Modeling

In our current work we chose to approximate the material behavior of our thrombus mimics as hyperelastic. Theoretically this means that the stress tensor was derived from a scalar function, the strain energy function. Practically, this implies that we ignore viscoelastic phenomena such as hysteresis, strain-rate dependence, stress-relaxation, etc. We chose this approach as it is commonly used as a first approximation to the mechanical behavior of soft tissues, while providing an appropriate framework for modeling materials that undergo very large deformations. As we planned on modeling our material as hyperelastic, we chose a quasistatic strain rate that renders the time-dependent phenomena negligible (i.e., we allow the material to equilibriate while it deforms). In future studies, we will investigate the time- and rate-dependent behavior of our thrombus mimics across various strain-rates and relaxation times. Of the three models, the Ogden model captured our mimics’ material behavior most accurately. It should also be noted that the Ogden model with only two parameters exceeded the quality of the Yeoh model with three parameters, further highlighting its ability to capture our mimics’ behavior. Thus, we recommend this model when using our data and suggest that other studies of the mechanical behavior of whole blood samples consider the Ogden model as well.

### 4.5. Mimic Composition and Blood Storage Time

We found that the mechanics of our thrombus mimics were relatively insensitive to coagulation conditions and blood storage time. To test the latter, we counted platelets and red blood cells across a period of 3 days and determined their concentrations. Those tests found little change in live cell concentration. Quantitatively, our numbers agree with those reported in the literature for bovine blood (Roland et al., 2014). Anecdotally, our findings also agree with statements by Chernysh et al., who state in their work that “RBCs are typically stored for up to 42 days, and we know from other studies that levels of ATP, 2,3-DPG fall very slowly over many days, and structural changes are also slow. On the other hand, platelets are only stored for a maximum of 5 days.” (Chernysh et al., 2020). Additionally, we investigated whether the toe-stiffness and/or calf-stiffness significantly varied over time and found no significant correlation there either. Thus, in agreement with Chernysh et al., we suggest that the mechanics of thrombus mimics like ours is stable between days 1 and 3 of storage.

### 4.6. Future Directions

This was the first step in our attempt of establishing an in-vitro thrombus mimic toward understanding thrombus mechanics. While we successfully generated and mechanically tested our thrombus mimics, explored their sensitivity to coagulation conditions, and identified their constitutive behavior, there is much work to be done. In the future, we will first compare our in-vitro thrombus mimics based on bovine blood to samples generated from human blood to ensure clinical relevance of our findings. Secondly, we will begin to generate mimics that resemble thrombi of various degrees of maturation. To this end, we will vary the mimics’ composition as well as co-polymerize collagen during clot generation. For validation of this approach, we will compare the histomechanics of our in-vitro thrombus mimics to samples excised from patients.

### 4.7. Limitations

Naturally, this work shows some limitations. First and foremost, we performed all studies on bovine blood, not human blood. Thus, care must be taken before extrapolating our findings to human thrombus. In terms of cell viability, our current manual method for tracking the number of live red blood cells and platelets, albeit standard and historically proven, is subject to high variability and thus uncertainty. Most importantly, our mimics were not formed in-vivo. While in-vitro formation ensures consistency, homogeneity, and conformation with geometric requirements, it produces “unnatural” thrombi. This implies, among other differences, that these mimics have not been invaded by inflammatory or synthetic cell species (Lee et al., 2015), demonstrate no neovascularization (Wakefield et al., 1999) or fissures or sinuous cavity formations (Fineschi et al., 2009), no fibrinolysis has occured (Longstaff and Kolev, 2015), no collagen has been deposited (Wakefield et al., 2008), no Lines of Zahn are apparent (Lee et al., 2012), and no shear flow-induced topical anisotropy is present (Campbell et al., 2010).

### 5. Conclusion

We developed a robust protocol to generate regularly-shaped, homogeneous thrombus mimics that lend themselves to simple shear testing under large deformation. Our data demonstrate that those mimics exhibited shear-stiffening, a negative Poynting effect, and hysteresis and that the material properties were insensitive to coagulation conditions and storage conditions (within the first three days). Additionally, we found that the Ogden material model fits these mechanical data excellently. Future studies will focus on comparing our findings to those of in-vivo thrombi and on extending our mimics to also represent maturing thrombus.

